# Genetic risk for neurodegenerative disorders, and its overlap with cognitive ability and physical function

**DOI:** 10.1101/219345

**Authors:** Saskia P. Hagenaars, Ratko Radakovic, Christopher Crockford, Chloe Fawns-Ritchie, International FTD-Genomics Consortium (IFGC), Sarah E. Harris, Catharine R. Gale, Ian J. Deary

## Abstract

**INTRODUCTION:** It is unclear whether polygenic risk for neurodegenerative disease is associated with cognitive performance and physical health.

**METHODS:** This study tested whether polygenic scores for Alzheimer’s disease (AD), Amyotrophic Lateral Sclerosis (ALS), or frontotemporal dementia (FTD) are associated with cognitive performance and physical health. Group-based analyses were performed to compare associations with cognitive and physical function outcomes in the top and bottom 10% for the three neurodegenerative polygenic risk scores.

**RESULTS:** Higher polygenic risk scores for AD, ALS, and FTD were associated with lower cognitive performance. Higher polygenic risk scores for FTD was also associated with increased forced expiratory volume in 1s and peak expiratory flow. A significant group difference was observed on the symbol digit substitution task between individuals with high polygenic risk for FTD and high polygenic risk for ALS.

**DISCUSSION:** Our results suggest overlap between polygenic risk for neurodegenerative disorders, cognitive function and physical health.

## 1. Introduction

Alzheimer’s disease (AD) is the most common form of dementia and it is expected that over one million people in the UK will be diagnosed with this disease by 2025 [1]. Pathological features of AD include amyloid B-plaques and neurofibrillary tangles [2], while degeneration of subcortical hippocampal regions and the medial temporal lobes is associated with the salient presenting memory impairment [3, 4].

Frontotemporal dementia (FTD), in contrast, is marked by degeneration of the frontal and anterior temporal lobes. Behavioural variant FTD (bvFTD) is the most common phenotype, and is marked by behaviour change and executive dysfunction (e.g., working memory, cognitive flexibility, response generation, and social cognition), with relative sparing of episodic memory and visuospatial skills [5].

The cognitive profiles of both AD and FTD are heterogeneous, in that impairments in executive functions, language, visuospatial skills, and memory have been recorded in both diseases [6]. Whereas evidence does exist that memory, visuospatial, and particularly language functions may be affected in bvFTD, the extent to which executive dysfunction explains these observations remains unclear [7, 8]. Adding to this complexity is the genetic, pathological, and neuropsychological overlap between FTD and Amyotrophic Lateral Sclerosis (ALS). ALS is a neurodegenerative disease marked by progressive loss of motor neurons in the brain and spinal cord, resulting in muscle wasting, spasticity, and death usually due to respiratory failure within three years [9]. Approximately 35% of ALS patients will present with mild cognitive and/or behavioural changes, of a similar nature to those seen in FTD, with an additional 15% meeting diagnostic criteria for both ALS and FTD [10]. Similarly, approximately 40% of FTD patients present with motor symptoms, and 15% meet ALS classification criteria [11].

Whereas the neuropsychological profiles of AD, ALS, and FTD can be heterogeneous, so too are the genetic profiles. The *APOE* e4 allele is the strongest genetic risk factor for late onset AD; however, numerous additional candidate and susceptibility loci have been established in recent years [12]. It has been suggested that late onset AD is due to susceptibility at multiple loci, and due to genetic and environmental interactions [13]. Early onset AD is autosomal dominant, accounting for approximately 1-5% of all cases, has been linked to variants in the *APP, PSEN1*, and *PSEN2* genes [14].

Autosomal dominant ALS accounts for approximately 5-10% of cases, with 85-90% of cases having no strong genetic linkage [15]. Familial ALS has been linked to the *SOD1, TARDBP, FUS*, and more recently *NEK1* genes [16, 17]. Conversely, up to 40% of FTD cases show a positive family history, with mutations in the *MAPT* and *GRN* genes being amongst the most common [18]. However, most frequently, the *C9ORF72* mutation has been associated with both familial ALS and FTD [19, 20]. The *C9ORF72* mutation can also be found in sporadic forms of ALS and FTD [21, 22], however the cause here is often unknown, and possibly due to environmental-genetic linkages [16]. A genome-wide association study (GWAS) of FTD [23] showed that the *C9ORF72* mutation mainly associates with the variant of FTD that overlaps with ALS, rather than the bvFTD variant, indicating genetic overlap between FTD and ALS.

Previous research has suggested a preclinical phase of reduced cognitive functioning for older adults who subsequently develop AD [24] and for those at genetic risk of developing AD [25-29]. Similar findings have been reported for individuals at risk of FTD [30, 31]. Given that not all those at genetic risk of developing AD or FTD go on to develop the disease, these findings may suggest that disease-specific carriers may be at risk of developing preclinical symptoms without necessarily developing the disease itself.

As previously noted, the genetic aetiology of AD, ALS, and FTD is multifactorial, and a significant proportion of genetic variance remains unexplained (missing heritability). As such, polygenic risk scores are becoming increasingly useful in the study of genetically complex diseases. Polygenic risk scores are valuable in aggregating genetic markers that on their own do not reach significance [32]. Recent research has explored the effects of disease-specific polygenic risk on cognitive performance. For example, a high polygenic risk of schizophrenia has been associated with a greater decline in cognitive functioning between childhood and older age [33], and reduced neural efficiency [34]. Similarly, high polygenic risk for autism has been associated with better cognitive ability in the general population [35, 36].

Higher polygenic risk for AD, based on genome-wide significant single nucleotide polymorphisms (SNPs) [37] and all common SNPs [36], has been associated with lower general cognitive ability and memory. Some research has suggested that healthy adults with a high polygenic risk for AD possess reduced brain cortical thickness [38]. To date however, no identifiable research has explored whether polygenic risk for FTD is associated with cognition, or whether high risk for ALS is related to cognitive, muscle, or respiratory function. As such, the aim of this study is to further explore whether polygenic risk for AD, FTD, or ALS is associated with cognitive performance, or with physical function measures known to be affected by motor neuron degeneration.

## 2. Materials and methods

### 2.1. Sample

This study includes baseline data and data from a web-based cognitive follow-up from the UK Biobank study, a large resource for identifying determinants of human diseases in middle aged and older healthy individuals. UK Biobank received ethical approval from the Research Ethics Committee (reference 11/NW/0382). This study has been completed under UK Biobank application 10279. A total 502,655 community-dwelling participants aged between 37 and 73 years were recruited between 2006 and 2010 in the United Kingdom, and underwent extensive testing including cognitive and physical assessments. All participants provided blood, urine and saliva samples for future analysis. For the present study, genomewide genotyping data was available for 112,151 individuals (58,914 females), aged between 40 and 70 years (mean age = 56.9 years, *SD* = 7.90) after the quality control process.

### 2.2. Measures

#### 2.2.1. Cognitive measures

Cognitive ability was measured using five different cognitive tests. These included tests of verbal-numerical reasoning (n = 36,035), reaction time (n = 111,484), memory (n = 112,067), trail making (part A: n = 23,822, part B: n= 23,812), and symbol digit substitution (n = 26,913). The verbal-numerical reasoning test consisted of a 13-item questionnaire assessing verbal and arithmetical deduction. Reaction time was measured using a computerized ‘Snap game’, during which participants were asked to press a button as quickly as possible when two cards on the screen were matching. Memory was measured using a pairs matching test, where participants were asked to memorize positions of matching pairs of cards, shown for 5s on a 3 by 4 grid. All cards were then placed face down and the participant had to identify the positions of the matching pairs as quickly as possible. The number of errors in this task was used as the (inverse) measure of memory ability. These tests have been previously described in more detail by Hagenaars et al. (2016)[36]. Executive functioning was measured using the trail making test parts A (TMT A) and B (TMT B), which were part of the follow up testing wave in UK Biobank, between 2014 and 2015. For the follow up testing, participants completed cognitive tests remotely via a web-based assessment. For TMT A, participants were instructed to connect numbers consecutively (which were quasi-randomly distributed on the touchscreen) as quickly as possible in ascending order by selecting the next number. TMT B is similar, but in this case letters and numbers had to be selected in alternating ascending order, e.g. 1 A 2 B 3 C etc. The difference between the raw scores for TMT A and TMT B was computed as TMT B minus TMT A (TMT B-A) [39]. Processing speed was measured using the symbol digit substitution test, which was also part of the follow up testing wave. Participants were shown a series of grids in which they were asked to match symbols to numbers according to a key presented on the screen. The score for processing speed was based on the number of correctly matched symbols. Participants who scored 0 or above 40 were removed from further analyses.

#### 2.2.2. Physical measures

Hand grip strength was measured for both the right and left hand, using a Jamar J00105 hydraulic hand dynamometer. Participants were seated upright in a chair with their forearms on the armrests, and were asked to squeeze the handle of the dynamometer as strongly as possible for three seconds. The maximum grip strength for each hand was measured in whole kilogram force units. This study used maximum grip strength based on the dominant hand, as indicated by the participant.

Lung function was assessed using a Vitalograph Pneumotract 6800 spirometer. Participants were asked to record two to three blows, lasting for at least 6 seconds, within a period of 6 minutes. The following outcomes measures were calculated by the computer: forced expiratory volume in 1s (FEV1), forced vital capacity (FVC), and peak expiratory flow (PEF). FEV1 is the amount of air, in litres, that is forcibly exhaled in 1 second following full inspiration. FVC is the amount of air, in litres, that is exhaled followed full inspiration. PEF is the maximum speed of exhalation following full inspiration. All three measures were standardized and individuals with a Z-score > 4 were excluded from FEV1 (n = 47), FVC (n = 73), and PEF (n = 28).

### 2.3. Genotyping and quality control

The interim release of UK Biobank included genotype data for 152,729 individuals, of whom 49,979 were genotyped using the UK BiLEVE array and 102,750 using the UK Biobank axiom array. These arrays have over 95% content in common. Quality control was performed by Affymetrix, the Wellcome Trust Centre for Human Genetics, and by the present authors; this included removal of participants based on missingness, relatedness, gender mismatch, and non-British ancestry. Further details have been published elsewhere [36, 40]. Variants with a minor allele frequency of less than 0.01 and non-autosomal variants were excluded from further analysis.

### 2.4. Statistical analysis

The UK Biobank genotyping data required recoding from numeric (1, 2) allele coding to standard ACGT format before being used in polygenic profile scoring analyses, this was done using a bespoke program [36]. Polygenic risk scores were created for AD [12], ALS [22], FTD, [23] in all genotyped participants using PRSice software [41]. Polygenic risk scores were also created for AD excluding SNPs within a 500 J kb window of apolipoprotein E (*APOE*) gene. PRSice calculates the sum of alleles associated with the phenotype of interest across many genetic loci, weighted by their effect sizes estimated from a GWAS of the corresponding phenotype in an independent sample. SNPs with a minor allele frequency < 0.01 were removed prior to creating the scores. Clumping was used to obtain SNPs in linkage disequilibrium with an r2 < 0.25 within a 250kb window. Five polygenic risk scores were then created for the three phenotypes containing SNPs selected according to the significance of their association with the phenotype, at thresholds of p < 0.01, p < 0.05, p < 0.1, p < 0.5, and all SNPs. The associations between the polygenic profiles and the target phenotype were examined in regression models, adjusting for age at measurement, sex, genotyping batch and array, assessment centre, and the first 10 genetic principal components to adjust for population stratification. The models including measures of lung function (FEV1, FVC, PEF) were also adjusted for smoking status and height, whereas the models for grip strength were additionally adjusted for height and weight. All models were corrected for multiple testing using the false discovery rate method [42].

Following calculation of each participant’s polygenic risk score at each of the above-mentioned thresholds, participants were classified in terms of their high or low polygenic risk for each neurodegenerative disease (AD, ALS and FTD). High risk was defined by the participant’s polygenic risk score falling within the top 10th percentile and low risk was defined by the participant’s polygenic risk score falling within the bottom 10th percentile for any of the neurodegenerative diseases. These participants were then grouped based on High Risk for AD (top 10th percentile for AD and bottom 10th percentiles for ALS and FTD), High Risk for ALS (top 10th percentile for ALS and bottom 10th percentiles for AD and FTD) and High Risk for FTD (top 10th percentile for FTD and bottom 10th percentiles for ALS and AD) to homogenise pure polygenic risk for each neurodegenerative disease in these groups. There were no overlapping participants in each of these groups. These High Risk for AD, High Risk for ALS and High Risk for FTD groups were compared on cognitive and physical variables (chosen based on significant polygenic profile-target phenotype regression models from the previous analysis). Shapiro-Wilk tests were used to assess normality of the data, following which group comparisons were performed either using Kruskal-Wallis H or ANOVA tests on cognitive and physical variables, applying a Holm-Bonferroni correction. Significant Kruskal-Wallis H or ANOVA results were followed up with post hoc tests (Tukey’s Honest Significant Differences or Mann Whitney U tests, respectively), where effect size was estimated using Cohen’s d.

## 3. Results

### 3.1. Polygenic risk analysis

The results for the polygenic risk analyses examining if polygenic risk for neurodegenerative diseases is associated with cognitive ability and physical health, using the best threshold (largest β), are shown in Table 1 and Figure 1. Full results including all five thresholds can be found in Supplementary Table 1. Polygenic risk scores for AD significantly predicted verbal-numerical reasoning (β = −0.023, *p* = 1.27 × 10^-5^) [36], memory (β = 0.011, *p* = 0.0001) [36], symbol digit substitution (β = −0.015, *p* = 0.0065), and TMT B (β = 0.017, *p* = 0.0047). Thus, individuals with higher polygenic risk for AD answered fewer verbal-numerical reasoning questions correctly, made more errors on the memory task, completed fewer items on the symbol digit substitution test, and took longer to complete TMT B. When excluding APOE from the AD polygenic risk score, the associations with symbol digit substitution attenuated to non-significance (Supplementary Table 1). Polygenic risk scores for ALS significantly predicted verbal-numerical reasoning (β = −0.019, *p* = 0.0004). Individuals with higher polygenic risk for ALS had lower scores on verbal-numerical reasoning. Polygenic risk scores for FTD significantly predicted TMT B (β = 0.017, *p* = 0.0041), FEV1 (β = 0.007, *p* = 0.0012), and PEF (β = 0.011, *p* = 4.77 × 10^-6^). Those with higher polygenic risk for FTD took longer to complete TMT B and had higher FEV1 and PEF scores. The associations between all scores at each threshold are shown in Supplementary Table 2.

**Table 1.** Associations between polygenic risk scores for Alzheimer’s disease (FDR p-value <= 0.018), amyotrophic lateral sclerosis (FDR p-value <= 0.0024), and frontotemporal dementia (FDR p-value <= 0.0041), and cognitive and physical measures controlling for age, sex, assessment centre, genotyping batch and array and 10 genetic principal components for population structure. pT, polygenic risk score threshold for best model; *, previously published by Hagenaars et al., (2016) [36]

|  | Alzheimer's disease |  |  | Amyotrophic lateral sclerosis |  |  | Frontotemporal dementia |  |  |
| --- | --- | --- | --- | --- | --- | --- | --- | --- | --- |
| | pT | $\beta$ | p | pT | $\beta$ | p | pT | $\beta$ | p |
| **Verbal-numerical reasoning** | 0.05 | -0.0229 | $1.27 \times 10^{-5*}$ | 0.05 | -0.0188 | **0.0004** | 1 | -0.0085 | 0.1036 |
| **Reaction time** | 0.5 | 0.0052 | 0.0700* | 1 | 0.0023 | 0.4315 | 0.5 | -0.0033 | 0.2489 |
| **Memory** | 0.1 | 0.0114 | **0.0001\*** | 0.1 | 0.0080 | 0.0074 | 0.05 | 0.0053 | 0.0737 |
| **Symbol digit substitution** | 0.5 | -0.0150 | **0.0065** | 0.01 | -0.0063 | 0.2613 | 0.01 | -0.0053 | 0.3316 |
| **Trail making part A** | 0.05 | 0.0144 | 0.0195 | 0.01 | 0.0087 | 0.1609 | 0.1 | 0.0104 | 0.0900 |
| **Trail making part B** | 0.5 | 0.0168 | **0.0047** | 0.1 | 0.0155 | 0.0101 | 0.1 | 0.0171 | **0.0041** |
| **Trail making part B-A** | 1 | 0.0104 | 0.0965 | 0.1 | 0.0166 | 0.0091 | 0.1 | 0.0093 | 0.1410 |
| **Grip strength** | 0.1 | -0.0016 | 0.3963 | 0.01 | 0.0032 | 0.0997 | 0.05 | 0.0020 | 0.2911 |
| **Forced expiratory volume in 1s** | 1 | -0.0027 | 0.2018 | 0.01 | 0.0022 | 0.3066 | 0.05 | 0.0068 | **0.0012** |
| **Peak expiratory flow** | 1 | -0.0039 | 0.1159 | 0.01 | 0.0017 | 0.4943 | 0.1 | 0.0113 | **$4.77 \times 10^{-6}$** |
| **Forced vital capacity** | 1 | -0.0014 | 0.4817 | 0.01 | 0.0019 | 0.3158 | 0.05 | 0.0027 | 0.1628 |

**Figure 1.**
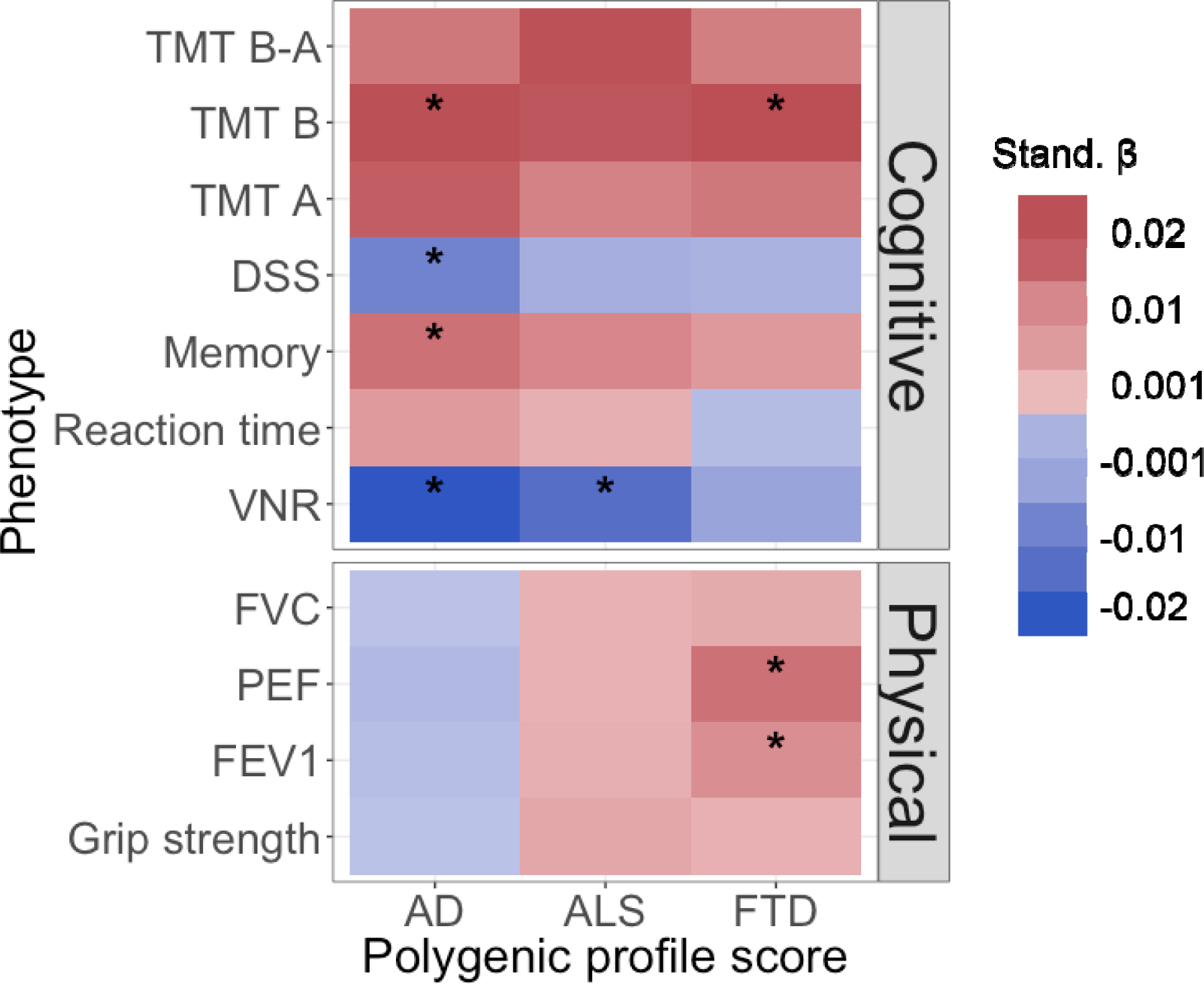
Heat map of associations between the polygenic profile scores for neurodegenerative disease and cognitive ability and physical health. Stronger associations are indicated by darker shades, red indicates a positive association, blue indicates a negative association. AD, Alzheimer’s disease; ALS, amyotrophic lateral sclerosis; FTD, frontotemporal dementia; TMT B-A, trail making part B - part A; TMT B, trail making part B; TMT A, trail making part A; DSS, digit symbol substitution; VNR, verbal numerical reasoning; FVC, forced vital capacity; PEF, peak expiratory flow; FEV1, forced expiratory volume in 1s. *, significant association after FDR correction (p-value <= 0.018 (AD), 0.024 (ALS), or 0.0041 (FTD)). Full results can be found in Supplementary Table 1.

**Table 2.**
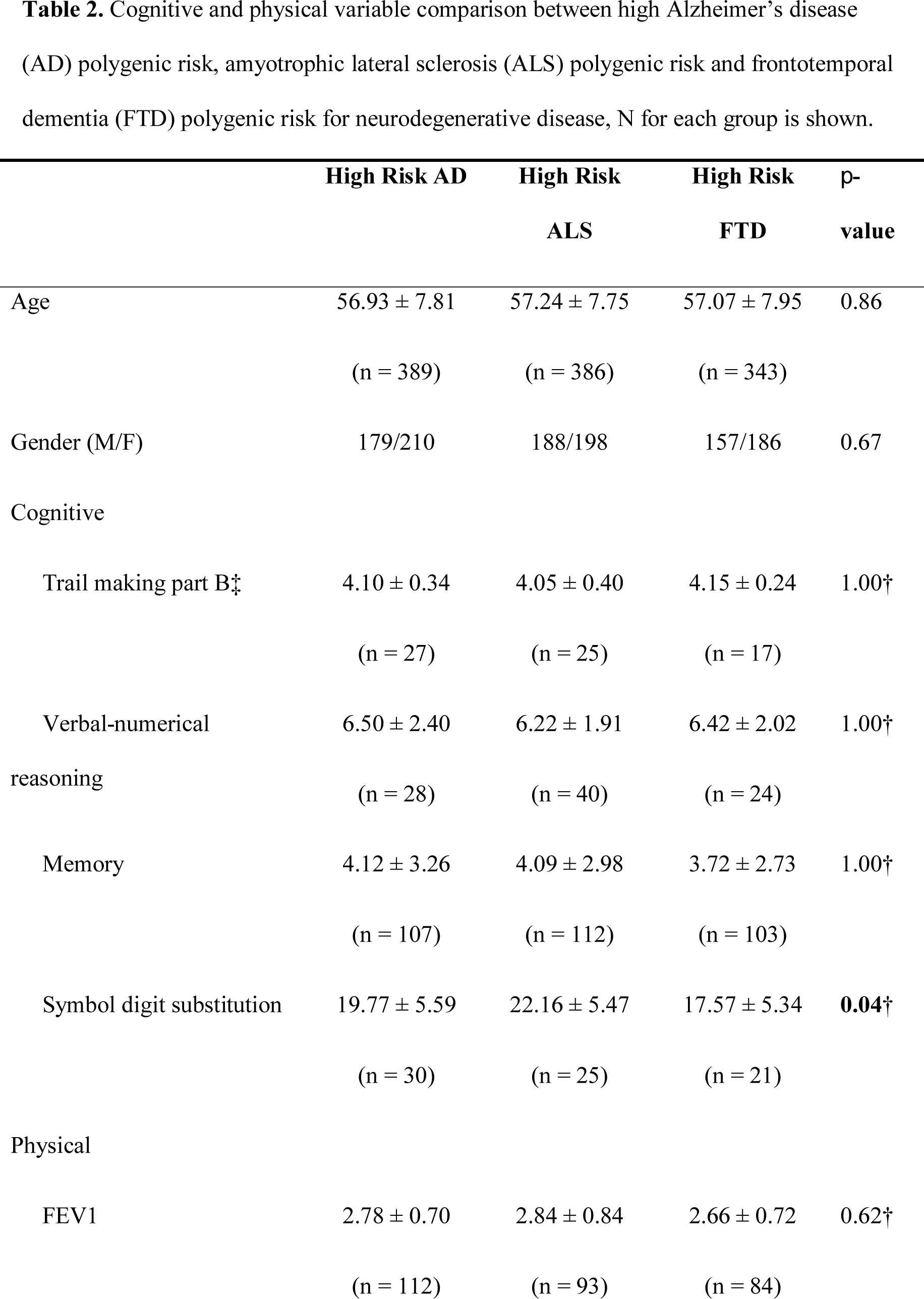
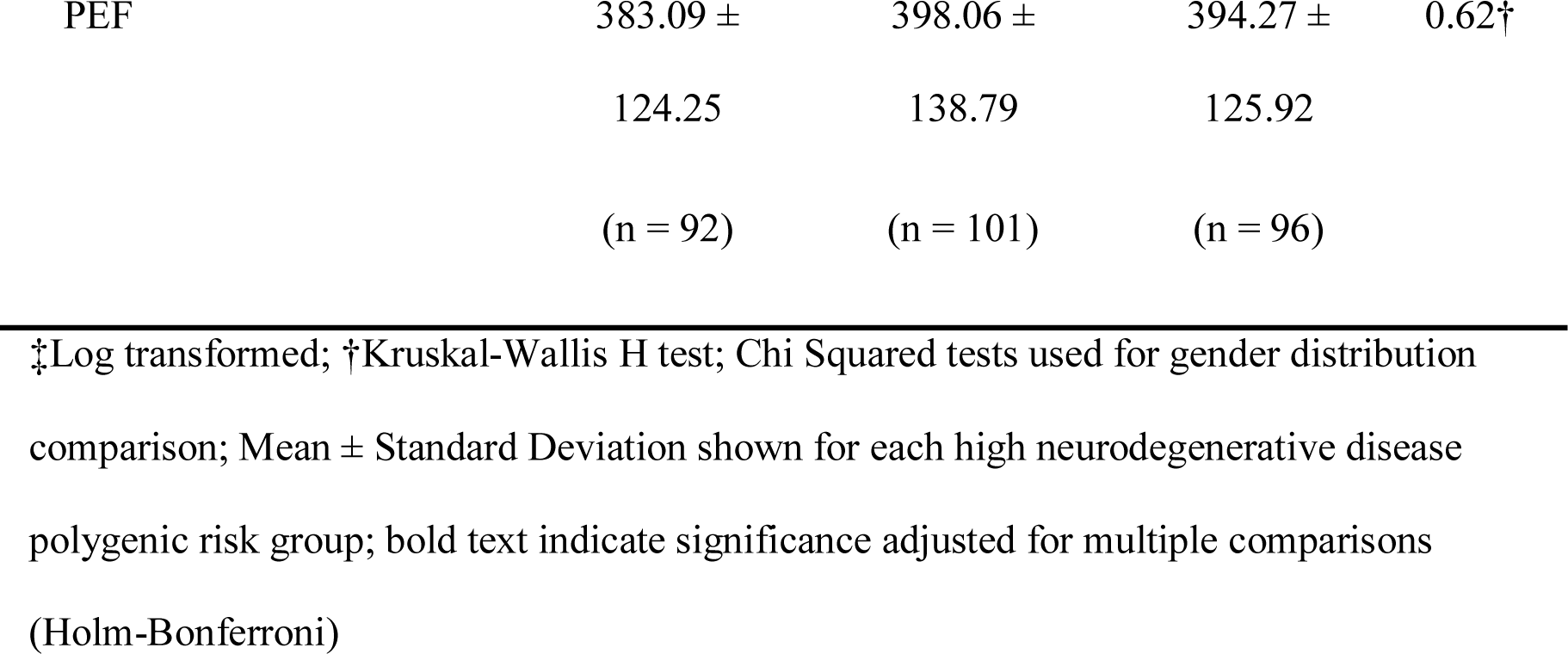
Cognitive and physical variable comparison between high Alzheimer’s disease (AD) polygenic risk, amyotrophic lateral sclerosis (ALS) polygenic risk and frontotemporal dementia (FTD) polygenic risk for neurodegenerative disease, N for each group is shown.

### 3.2. Polygenic Inter-group Comparison

Table 2 shows the comparison of polygenic risk groups on cognitive and physical variables. There was no significant difference between those categorised as having high polygenic risk for AD, ALS, or FTD on age or gender distribution. The only significant difference between these groups was observed with one of the cognitive variables, specifically, symbol digit substitution task. Follow up, post hoc analysis showed that participants with a high polygenic risk for FTD performed significantly worse on the symbol digit substitution task when compared to participants with a high polygenic risk for ALS (U = 396, *p* < 0.01, *d* = 0.85). No significant differences were observed between other groups on cognitive and physical variables.

## 4. Discussion

The present study aimed to explore whether polygenic risk for AD, FTD, or ALS is associated with cognitive performance, grip strength, or lung function measures. Using the large cognitive, physical and genotypic data available in the UK Biobank, in concordance with the most up to date GWAS consortia of neurodegenerative disease (AD, ALS and FTD), our results showed a novel relationship between cognitive and physical function variables and polygenic risk for neurodegenerative conditions in a healthy population, particularly in the case of AD.

The findings of this study demonstrate that higher polygenic risk of AD is associated with numerous cognitive functions; specifically, reduced performance in verbal-numerical reasoning, memory, processing speed (symbol digit substitution), and executive functioning (TMT B). Sensitivity analysis indicated that the association between AD and cognitive ability (except symbol digit substitution) was not driving by the *APOE* gene. Although an association between polygenic risk of AD and cognitive ability - that is significant associations with verbal-numeric reasoning and memory - in UK Biobank, has been reported previously [36], the present study extends these findings to measures of executive functioning and processing speed. Unlike AD, higher polygenic risk for ALS demonstrated only one significant relationship with cognitive function, namely verbal-numerical reasoning. Higher polygenic risk for FTD was only associated with reduced performance on the TMT B.

Physical function measures used in this study were chosen based on their relevance in clinical cases of FTD and ALS, in particular, measures of grip strength and lung function. Counterintuitively higher risk of FTD was related to better respiratory functioning (FEV1 and PEF). Finally, a higher risk of ALS was not associated with grip strength or measures of lung function. It is worth noting, however, that the amount of variance explained by the polygenic risk scores was very small (< 0.01%). As such, these finding may be spurious.

In a small subset of participants, those with a high polygenic risk for FTD performed worse on a processing speed task (symbol digit substitution task) compared to those with high polygenic risk for ALS. There has been an emphasis on linkage and continuum-based relationship between ALS and FTD, this finding based on pure genetic risk could provide insight towards sub-clinical cognitive impairment. The cognitive profile of ALS largely mirrors that of FTD, albeit in a milder form. Patients with FTD have demonstrated processing speed impairments, both cross-sectionally and longitudinally [43, 44]. However, studies of cognition in ALS have largely ignored tasks of processing speed due to the confounding presence of motor impairment in this patient group. That said, current research suggests unaffected processing speed in patients with ALS. As such, previous research combined with the findings herein may suggest a future avenue for detecting FTD syndromes in patients with ALS. However, further studies are needed to explore the reliability of differences between polygenic risk of FTD and ALS, and between people diagnosed with these diseases.

There are several limitations to note regarding the present study. While the cognitive tasks in the present study covered a number of domains, the measures are brief and non-standardized. As such, the sensitivity to detect small differences in cognitive ability are limited. Additionally, due to the nature of self-administration on a computer (in case of the measures for the symbol digit substitution test and the trail making test), the environment under which the tasks are performed could not be fully standardised.

Furthermore, while it is interesting to speculate how the findings of the present study may relate to those individuals who actually have AD, FTD, or ALS, it is important to remember that the participants of this study were healthy individuals. Those described as high polygenic risk for a particular disease are only at higher risk when compared to other participants of this study. It would be incorrect to describe these individuals as being at a high risk of developing a neurodegenerative disease more generally.

Future research may explore whether these findings replicate given more extensive and controlled measures of cognitive functioning. Provided that FTD is primarily a disease marked by changes in behaviour, the inclusion of such measures would be informative. Additionally, as previous research has suggested that reduction of cortical thickness was associated with polygenic risk for AD in healthy adults [38], it would be informative to further explore the neural correlates of polygenic risk for different neurodegenerative disorders (e.g. ALS and FTD) in this sample.

## 5. Conclusion

The present study confirmed and extended previous findings that polygenic risk for AD is associated with multi-domain cognitive functioning in healthy adults. Additionally, the findings of this study demonstrate that polygenic risk for ALS is associated with verbal-numeric reasoning, while polygenic risk for FTD was associated with executive functioning. Physical function measures commonly affected in patients with ALS, were not associated with polygenic risk of ALS in healthy adults. However, higher polygenic risk significantly predicted better lung function.

## Acknowledgments

This research was conducted using the UK Biobank Resource. The work was undertaken in The University of Edinburgh Centre for Cognitive Ageing and Cognitive Epidemiology, part of the cross council Lifelong Health and Wellbeing Initiative (MR/K026992/1); funding from the BBSRC and Medical Research Council (MRC) is gratefully acknowledged. This report represents independent research part-funded by the National Institute for Health Research (NIHR) Biomedical Research Centre at South London and Maudsley NHS Foundation Trust and King’s College London. CC is supported by the Euan MacDonald Centre for Motor Neurone Disease Research. CF-R is supported by Dementias Platform UK (DPUK), funded through the MRC (MR/L023784/2). The authors would like to thank the Project MinE GWAS Consortium. The authors thank the International FTD-Genomics Consortium (IFGC) for providing phase I summary statistics data for these analyses. Further acknowledgments for IFGC, e.g. full members list and affiliations, are found in the online supplementary files.

## Collaborators

R Ferrari; D G Hernandez; M A Nalls; J D Rohrer; A Ramasamy; J B J Kwok; C Dobson-Stone; P R Schofield; G M Halliday; J R Hodges; O Piguet; L Bartley; E Thompson; E Haan; I Hernández; A Ruiz; M Boada; B Borroni; A Padovani; C Cruchaga; N J Cairns; L Benussi; G Binetti; R Ghidoni; G Forloni; D Albani; D Galimberti; C Fenoglio; M Serpente; E Scarpini; J Clarimón; A Lleó; R Blesa; M Landqvist Waldö; K Nilsson; C Nilsson; I R A Mackenzie; G-Y R Hsiung; D M A Mann; J Grafman; C M Morris; J Attems; T D Griffiths; I G McKeith; A J Thomas; P Pietrini; E D Huey; E M Wassermann; A Baborie; E Jaros; M C Tierney; P Pastor; C Razquin; S Ortega-Cubero; E Alonso; R Perneczky; J Diehl-Schmid; P Alexopoulos; A Kurz; I Rainero; E Rubino; L Pinessi; E Rogaeva; P St George-Hyslop; G Rossi; F Tagliavini; G Giaccone; J B Rowe; J C M Schlachetzki; J Uphill; J Collinge; S Mead; A Danek; V M Van Deerlin; M Grossman; J Q Trojanowski; J van der Zee; M Cruts; C Van Broeckhoven; S F Cappa; I Leber; D Hannequin; V Golfier; M Vercelletto; A Brice; B Nacmias; S Sorbi; S Bagnoli; I Piaceri; J E Nielsen; L E Hjermind; M Riemenschneider; M Mayhaus; B Ibach; G Gasparoni; S Pichler; W Gu; M N Rossor; N C Fox; J D Warren; M G Spillantini; H R Morris; P Rizzu; P Heutink; J S Snowden; S Rollinson; A Richardson; A Gerhard; A C Bruni; R Maletta; F Frangipane; C Cupidi; L Bernardi; M Anfossi; M Gallo; M E Conidi; N Smirne; R Rademakers; M Baker; D W Dickson; N R Graff-Radford; R C Petersen; D Knopman; K A Josephs; B F Boeve; J E Parisi; W W Seeley; B L Miller; A M Karydas; H Rosen; J C van Swieten; E G P Dopper; H Seelaar; Y A L Pijnenburg; P Scheltens; G Logroscino; R Capozzo; V Novelli; A A Puca; M Franceschi; A Postiglione; G Milan; P Sorrentino; M Kristiansen; H-H Chiang; C Graff; F Pasquier; A Rollin; V Deramecourt; T Lebouvier; D Kapogiannis; L Ferrucci; S Pickering-Brown; A B Singleton; J Hardy; P Momeni

